# 3’ RNA sequencing does not increase power or reduce costs for gene expression analysis

**DOI:** 10.1101/2022.04.13.488225

**Authors:** Taylor M Crow, JA Gill, Andrew Whitehead, Daniel E Runcie

**Affiliations:** Department of Plant Sciences, University of California Davis, Davis, USA; Department of Environmental Toxicology,, University of California Davis, Davis, USA

**Keywords:** sample, article, author

## Abstract

**Background:** Sequencing RNA transcripts for gene expression profiling is a popular and important technique with broad utility in biological sciences. We set out to comprehensively compare the two most popular methods for generating sequencing libraries for differential gene expression analysis: 3-end sequencing, which generates libraries from the 3’ end of an RNA transcript; and traditional RNA sequencing, which generates libraries from whole RNA transcripts. We include three species in our experiment to test whether our findings replicate across genomes and genome assemblies.

**Results:** We found similar levels of precision and power to detect differentially expressed genes between the two methods. Notably, whole transcript RNA-seq performed better in the non-traditional model species included in our study.

**Conclusion:** Overall, we recommended whole transcript RNA sequencing for the added benefits of alternative splicing detection, and gene-body variant detection.

## Introduction

High through-put RNA sequencing is widely used in biological research as a powerful tool for quantifying gene expression. The transcriptome reflects the state of gene expression at a cellular level, and is highly dynamic and responsive to external perturbations, which makes it an ideal quantitative molecular phenotype for a variety of biological questions. For example, how does expression change in response to environmental stressors? What cellular processes are important for acclimating to seasonal weather patterns? What genes are involved in the shift between vegetative and reproductive growth? Many basic and applied research hypotheses can be tested and validated using RNAseq, however, many questions and assumptions about RNAseq have not be adequately explored. One of the first questions a researcher needs to answer is: what type of RNAseq library construction should I choose?

The first step in RNAseq library construction is converting mRNA into cDNA libraries compatible for high-throughput sequencing platforms. Reverse transcription is primed with the addition of either random single-stranded hexamers–hybridizing along the length of the mRNA molecule–or through oligo d(T) primers–targeting the 3’ polyadenylated tail of the mRNA. This step is called first strand synthesis, where the single-stranded mRNA molecule is paired with a complimentary DNA (cDNA) strand. For the purposes of this manuscript, we refer to the two priming methods as whole transcript (WT) for random primers, and 3’ for libraries constructed with an oligo d(T) primer.

While both WT and 3’ libraries can accurately identify actively expressed genes (using strand-specific priming methods), their outputs permit and require different downstream applications. WT libraries sequence across the full-length of a transcript, making it possible to quantify alternative splicing of transcripts at a single locus. 3’ libraries can only distinguish among transcript families that have different 3’UTRs. However, expression-level estimates from WT libraries show a transcript-length bias where longer transcripts are sequenced at a higher read count relative to their expression. This can lead to biases for downstream analyses, e.g., Gene Ontology (GO) enrichment that need a length-bias correction [1]. 3’ libraries do not have a transcript-length bias because priming with a poly-A primer guarantees that only a single library molecule is made per transcript molecule.

Various authors have reasoned that either WT or 3’ libraries are inherently superior for differential gene expression analysis, based on arguments of cost, statistical power, and ease of down-stream analysis [2, 3, 4, 5]. Previous studies have found conflicting results when testing the difference in power between WT and 3’ (Table 1). For example, Tandonnet and Torres 2017 tested two treatments in a non-traditional model insect species (New World screw-worm fly) and detected 150 more DE contigs with 3’ than with WT [4]. However, they were unable to confidently annotate most of these contigs to gene models because they arose from 3’ UTR regions. The authors suggested that 3’ may have more power than WT for the same sequencing depth/cost but would be most appropriate for organisms with high-quality reference genomes. Xiong et al. 2017 tested three treatments (including untreated control) in a human commercial cell line and detected 15 percent more genes with WT libraries than 3’ libraries [3]. However they stated that WT libraries were more expensive and so experiments using 3’ could afford greater numbers of biological replicates. Ma et al. 2019 tested two treatments in a single system and detected between 58-72 percent more DE genes with WT libraries than 3’ libraries depending on read depth [5]. While WT libraries did suffer from a length bias, they stated that WT libraries were likely better for non-model organisms.

**Table 1:**
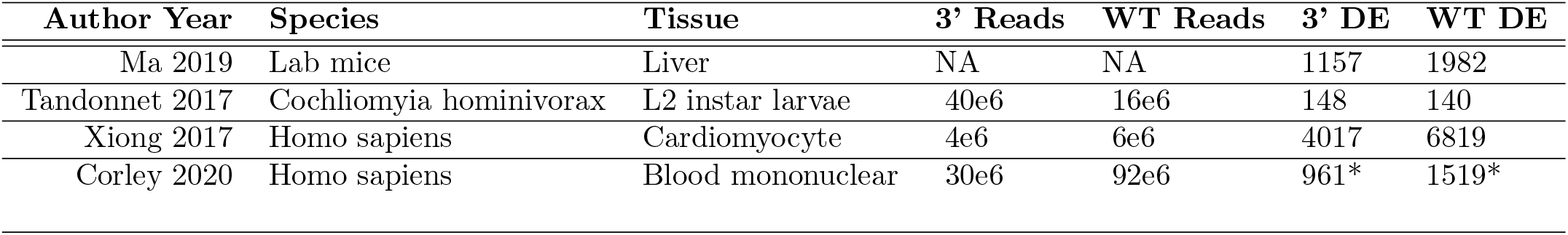
Summary of previous peer-reviewed research explicitly comparing RNA-seq priming methods. *These differentially expressed gene numbers are from the subset of data using salmon

We propose that many of the arguments in these papers are based on misconceptions about the relative advantages of the two priming methods for RNAseq:Library construction costs: Several authors have stated that 3’ libraries are less expensive [2, 3]. 2) Sequencing costs: authors have argued that 3’ libraries can be sequenced to lower depth without sacrificing precision or statistical power, thus reducing costs [2].

Neither of these issues actually favors 3’ libraries in general. While price differences certainly exist between commercial library construction kits, the difference between WT and 3’ library protocols is the type of oligo used to initiate reverse transcription, which has little to no difference in material costs [6].

The issue of sequencing costs differences between WT and 3’ libraries is a more subtle topic. Sequencing mRNA using WT can result in multiple library molecules primed and sequenced from the same mRNA molecule. Therefore, it is possible that the same mRNA molecule will be counted multiple times (e.g. two fragments from the same mRNA are sequenced). The two reads from the same mRNA molecule do not provide independent estimates of the expression of the corresponding transcript, so the second read is useless for quantification and a waste of resources. If quantification software is unable to determine that the second read is actually from the same mRNA molecule as the first and double-counts the molecule, this is effectively pseudoreplication and will lead to elevated false positive rates under models that assume a Poisson process (e.g. *edgeR, DESeq2*). The 3’ procedure is not subject to this issue because only a single library molecule can be reverse transcribed per transcript so no reads are wasted; every new read counts a new mRNA molecule.

However, this “double-counting” of mRNA molecules is actually extremely uncommon in most WT experiments, unless the input RNA quantities are exceptionally small, and sequencing depth is exceptionally high. For example, a typical RNAseq library preparation protocol recommends starting with approximately 50ng of mRNA. If the average gene length of a study organism is 2kb, such as in Eukaryotes [7], then a single sample would contain 46 billion mRNA molecules (The average molecular weight of a RNA base is 321.47 g/mol, so an average transcript will have a molecular weight of 643 Kg/mol. 50ng / 643Kg/mol * 6.022e23 = 46 billion molecules, calculations from: https://nebiocalculator.neb.com/#!/ssrnaamt, accessed September 1, 2021). RNAseq libraries are generally sequenced to a depth of 1-20 million reads. Since 3’ libraries sequence either 0 or 1 read per mRNA molecule, with 20 million sequenced reads, *≈* 0.04% of the original mRNA molecules will be sequenced. With WT libraries, if mRNA molecules are broken into 300nt fragments, there will be approximately 300 billion fragments, of which *≈* 0.007% will be sequenced. For any given mRNA molecule that has one read sequenced, the chance of a second being sequenced is also *≈* 0.007% for an mRNA molecule broken into 2 fragments, or up to *≈* 0.07% for an mRNA molecule broken into 11 fragments. This may well happen a few times in an RNAseq sample, but the vast majority of mRNA molecules will be represented by only a single sequenced read in WT libraries. Therefore, the effect of pseudoreplication in WT RNAseq is extremely low, and unlikely to affect analytical pipelines.

It is true that longer transcripts will have a higher probability of being counted (at least 1 time) than shorter transcripts in a WT library, meaning that longer transcripts will be measured with more precision by WT libraries and shorter transcripts will be measured with more precision with 3’ libraries. For the shorter transcripts, 3’ libraries could be sequenced to lower depth than WT libraries and achieve equal precision. For example, a transcript of 50% the average transcript length (formally, a weighted average where transcript lengths are weighted by their expression) could be measured equally by a 3’ library sequenced to 50% the depth of an equivalent WT library. However this would come at the cost of much less precision for longer transcripts in the same library. A transcript of 2x the average transcript length would then receive only 25% of the read counts as the equivalent WT library. Also, the lack of length-bias does not mean that 3’ libraries measure each transcript with equal precision – higher expressed transcripts will still be measured with more precision by either protocol, and transcripts vary by many orders of magnitude in expression level but only by several fold in length, meaning that the contribution to heteroskedasticity of transcript length in WT libraries is not especially important relative to other factors.

While the analytical results above suggest that WT should not have higher costs or lower statistical power than 3’ libraries, we aimed to test this result experimentally. One possible confounding factor could be the relative abilities to correctly map and annotate RNAseq reads from each method, which depends on genome size and annotation quality, and varies among species. We therefore designed a study to carefully measure the differences in statistical power between experiments using WT vs 3’ libraries across three species with qualitatively different genomic and transcriptomic resources. Notably, unlike previous studies comparing WT and 3’ sequencing methods, we isolated the technical variation inherent to the RNAseq measurement process from the biological variation in order to make the comparisons more precise. We find no basis in the common assumption that WT libraries generate pseudoreplication of fragments, increasing type 1 error. Furthermore, 3’ libraries do not suffer from the same length bias as WT libraries which are more likely to over-represent longer transcripts. Finally, we find a small advantage in WT library’s ability to detect DE genes, most prominent in our non-traditional model system, suggesting WT is a better option for non-traditional model organisms that are perhaps lacking in genome assembly and annotation quality and completeness compared to more traditional model species.

## Materials and Methods

### Experimental design

We designed an experiment to quantify the effect of each library priming method on our ability to detect differentially expressed genes in 3 species. In the simplest experiment with two treatment groups, differential expression is tested by comparing a t-statistic, calculated as the ratio between an estimated change in expression divided by its standard error, to a null distribution. The library priming method could affect the power to detect differentially expressed genes by altering either the numerator (estimated effect size) or the denominator (within-group variance) of this ratio. If effect sizes are measured as log-fold-changes, it is unlikely that the two priming methods will differ in the estimated effect sizes (beyond sampling error) because longer genes should have proportionally more reads in both treatment groups with WT libraries. However, if pseudoreplication occurs in WT libraries, the within-group variance may be larger, leading to lower power. Therefore, our experiment was designed to compare the within-group variances of expression estimates between WT and 3’ libraries.

In a typical RNAseq experiment, biological replicates of each treatment group are used to test the effect of a treatment on expression by measuring the within-group variance. Previous experimental comparisons of WT and 3’ libraries have compared the results of experiments where some biological replicates are measured with WT libraries and others with 3’ libraries [2, 3, 4, 5]. However, in such experiments, testing whether the within-group variation (denominator of the t-statistics) is greater for WT than for 3’ libraries is under-powered because within-group variance is composed of both biological variation in mRNA levels among samples and measurement error in estimating mRNA levels inherent to the RNAseq process itself (including Poisson sampling error, any random error introduced during the library preparation process itself, and any random error in read mapping or counting).

We therefore used an experimental design to directly compare the magnitudes of measurement error inherent to WT and 3’ libraries. Rather than collecting multiple biological replicates from two treatment groups, we created a single sample from each of two treatments by pooling multiple samples together to get sufficient mRNA. We then split each pool into multiple identical RNA samples and used each to build either a WT or a 3’ library (Figure 1). Therefore, all variation in expression estimates from libraries created from the same pool is entirely technical in origin and we can directly compare the magnitudes of the among-sample (within-pool) variance for each gene between libraries made with each priming method.

**Figure 1:**
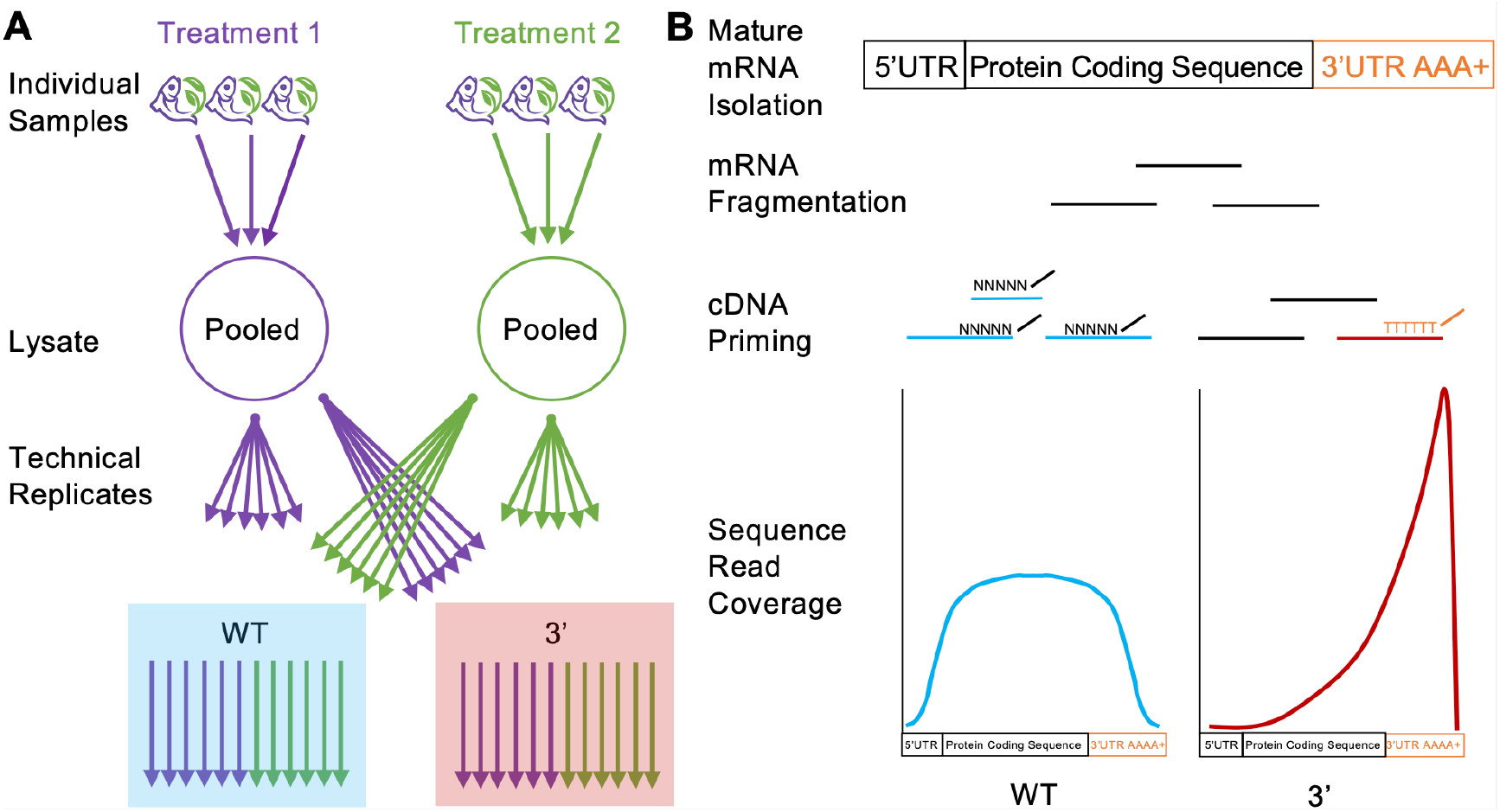
Graphical representation of experimental design to compare the magnitudes of technical noise in 3’ and WT libraries. **A**) 3 biological replicates per species were combined into a common pool then split into 6 equimolar technical replicates per library construction method (WT or 3’). **B**) Each technical replicate was individually fragmented by high temperature (95°C) and magnesium (Mg) ions and then first strand cDNA was synthesized either with poly-A (3’) or random hexamer (WT) primers. 3’ libraries are therefore 3’ biased and generate sequence only in the 3’ UTR or 3’ CDS. WT libraries are less biased along the mRNA molecule.

Below, we describe the detection of “significant” differences in expression between the two biological samples of each species as “differentially expressed genes”, or DEG. Note that we are not claiming that these lists of genes are biologically interesting in any sense outside of this experiment because the treatments applied to each species are not biologically replicated. Nevertheless, the analysis of differences between the two groups of technical replicates is exactly the same as if they had been biological replicates, and since all genes will differ to some extent in expression between the two pools, we can use this design to evaluate the effect of priming method on statistical power.

### Biological samples

Three species were selected for testing RNAseq library priming methods: Arabidopsis *Arabidopsis thaliana*, Maize (*Zea mays* spp. *mays*), and Pacific herring (*Clupea pallasii*). We used genetically homozygous genotypes of Arabidopsis (Col-0) and Maize (B73), and samples from a single wild population (Puget Sound, WA) of Pacific herring. The treatment groups for each species were as follows: Arabidopsis samples were collected at two different times of day; Maize samples were taken from two different growing temperatures; and Pacific herring samples were immediately late-stage (pre-hatch) embryos that had been exposed to crude oil and control (no oil) treatments during embryogenesis. Male and female adult Pacific herring gonads were collected from Puget Sound, WA by the Washington Department of Fish and Wildlife in 2017, and gametes were combined in the lab to form embryos for exposure experiments.

### mRNA extraction and creating technical replicates

All samples were stored at −80 °C. Tissue samples were homogenized using stainless steel beads in the SPEX Geno/Grinder (Metuchen, NJ, USA) at 1200 RPM for 1 minute. Pacific herring samples required a second round of homogenization at the same settings. Ground lysates were suspended in 200ul of Lysis binding buffer [6], incubated at room temperature for 10 minutes, centrifuged for 10 minutes at 13,000rpm, and the supernatant, referred to as cleared lysate, was retained.

The cleared lysate for each treatment group was combined to create a single mRNA pool with sufficient mRNA content to be subsequently divided into 6 200*µ*l identical samples of mRNA to create 6 technical replicates per treatment group. mRNA was extracted separately from each technical replicate using oligo (dT)_25_ beads (DYNABEADS direct™) to enrich for polyadenylated mRNA.

### RNA-seq library prep

We prepared strand specific RNA-seq libraries using the BRaD-seq protocol [6], and used 14 cycles of PCR at the enrichment phase. Library preparations for all species and for both priming methods were identical, except for the primer used for first strand cDNA synthesis. In this study we use a 3’ priming method (3’) to refer to libraries made with an oligo dT primer–which primes mRNA transcripts from the 3’ end–and whole transcript (WT) to refer to libraries made using a random hexamer to prime fragments from along the length of the transcript. For 3’ libraries, we used the primer: GTGACTGGAGTTCAGACGTGTGCTCTTC-CGATCTTTTTTTTTTTTTTTTTTTT, and for WT libraries, we used GTGACTGGAGTTCAGACGTGTGCTCTTCCGATCTNNNNNNNN, where N represents a random nucleotide as suggested in [6]. Finished libraries were quantified using the Quant-iT™ PicoGreen dsDNA high sensitivity kit, and normalized to 1ng/ul. We took 2ng per library and multiplexed the 36 samples for sequencing. The libraries were sequenced in 2 lanes of an Illumina HiSeq X platform, generating a mean of 6.1 million paired reads per sample. Raw reads were quality checked with FastQC v.0.11.5 [8]. Low quality reads (q*<*20), adapters, and reads less than 25-bp were removed using Trimmomatic v.0.36 [9].

### Genomic resources

Assembled reference genomes, and gene transfer format (GTF) files were downloaded from NCBI (https://www.ncbi.nlm.nih.gov/sra) [10]. We used the Arabidopsis accession number *GCF* 000001735.4 version *TAIR*10.1; the maize accession number *GCF* 000005005.2 version *B*73 *Ref Gen v*4. Pacific herring does not yet have an assembled genome, therefore we used the reference genome from a closely related species, *Clupea harengus* (Atlantic herring), accession number *GCA* 900700415.1 version *Ch v*2.0.2. The genomes varied in size and assembly and annotation quality. Arabidopsis has the highest quality assembly and annotation and smallest genome, maize has the largest genome, and herring has the lowest quality assembly and annotation (Table 2).

**Table 2:** Summary data of reference genome quality for each species used in this study. Source of the reference genome is NCBI (ncbi.nlm.nih.gov*\*sra).

|  | Metric | Atlantic herring | Arabidopsis | Maize |
| --- | --- | --- | --- | --- |
| 1 | Scaffold N50 | 1,842,407 | 23,459,830 | 10,679,170 |
| 2 | Contig N50 | 21,338 | 11,194,537 | 1,279,870 |
| 3 | Percent Complete | 84.7 | n/a | n/a |
| 4 | Genome Coverage | 72.0x | n/a | 65.0x |
| 5 | Total size | 808,340,743 | 119,668,634 | 2,134,373,047 |
| 6 | Release date | 2019 | 2015 | 2017 |

### RNA read mapping, quantification

Trimmed reads were mapped to reference genomes with Hisat2 [11] using default settings. The *featureCounts* function from Subread [12] was used to count transcript features from all samples, and sum the counts to the gene level.

### Differential expression pipeline

We developed a pipeline to detect differential expression that would account for variation in read depth across individuals, and bootstrapped the analyses to determine whether the results were consistent. Our pipeline was designed to reflect common experimental methods used in RNA-seq projects.

Samples with low total read counts (less than 100k reads), were removed from analysis. Since the remaining samples still varied in read counts, we randomly downsampled without replacement to the sample with the lowest number of mapped reads. All analysis were repeated 5 times with different downsampled datasets.

After down-sampling, WT and 3’ libraries of each species were processed independently through our differential expression pipeline. The downsampled gene count data were normalized using the weighted trimmed mean of M-values (TMM), using the *calcNormFactors* function in *edgeR* [13]. The *voom* function [14] in the *limma* package [15] was used to convert read counts to log2-counts per million (log2CPM), and estimate mean-variance trends to account for heteroscedasticity when testing for differential expression between the mRNA pools. The *lmFit* function from *limma* was used to fit a model using the pool ID as a predictor variable. The *ebayes* function was used to preform an empirical bayes shrinkage, and estimate an adjusted P-value after correcting for multiple testing using Benjamini-Hochberg correction.

## Results

We sequenced the multiplexed library on two lanes of Illumina HiSeq X platform using 150 bp paired-end reads. Previous work has shown that read length (50, 100, 150 bp) has no effect on the detection of differentially expressed genes [16]. The mean number of reads, trimming results, and mapping rate to the reference genome varied by species and library prep method (Table 3). Overall, 3’ libraries had more raw reads prior to trimming in Arabidopsis and Maize, but fewer in Pacific Herring. Since this effect would necessarily go away if all libraries were of the same type (because the total number of reads per lane is constant), we excluded variation in raw read counts in downstream analysis by down-sampling all libraries to the same total read number. For all three species, an average of 27% 3’ raw reads were trimmed (removed due to low quality), while 7% were trimmed in WT. Expression quantified using *featureCounts* resulted in approximately 15k, 18k, and 8k genes with greater than 5 reads in more than half of individuals identified in maize, arabidopsis, and Pacific herring respectively (Table 4).

**Table 3:** Sequencing results for the number of raw, processed, and percent of uniquely mapped reads for each RNA-seq method and sample condition.

|  | Species | Priming | Raw reads | Trimmed | Reads remain | Mapping |
| --- | --- | --- | --- | --- | --- | --- |
| 1 | arabidopsis | 3' | 10187619.50 | 24.00 | 7708045.25 | 98.00 |
| 2 | arabidopsis | WT | 6095645.45 | 7.00 | 5733226.55 | 99.00 |
| 3 | herring | 3' | 3598875.50 | 27.00 | 2588210.75 | 95.00 |
| 4 | herring | WT | 3798496.20 | 7.00 | 3562483.50 | 90.00 |
| 5 | maize | 3' | 17585885.42 | 31.00 | 11992257.42 | 93.00 |
| 6 | maize | WT | 11342424.25 | 7.00 | 10628689.75 | 96.00 |

**Table 4:** Summary of results from analysis for each species and primer. Total number of analyzable genes (genes with 5 or more reads in half of individuals), DE = total number of differentially expressed genes and %DE = the percent of total differentially expressed genes divided by the total number of analyzable genes. Differential expression threshold was genes below an FDR level of 0.05.

|  | Species | Primer | Total | DE | DE (%) |
| --- | --- | --- | --- | --- | --- |
| 1 | arabidopsis | 3' | 18688 | 3398 | 18.18 |
| 2 | arabidopsis | WT | 18142 | 3459 | 19.06 |
| 3 | herring | 3' | 7783 | 388 | 4.98 |
| 4 | herring | WT | 9035 | 811 | 8.98 |
| 5 | maize | 3' | 15410 | 5373 | 34.87 |
| 6 | maize | WT | 15361 | 5814 | 37.85 |

To explore if WT or 3’ libraries differed significantly in either estimates of treatment effects, or precision (within-group standard errors of estimated expression), we first assessed whether the estimates of treatment effects differed between WT and 3’ libraries. The difference in estimated logFC was not biased above or below zero for either low or high expressed genes (Figure 2A), or for short of long genes (Figure 2B). We next assessed whether the standard error of effect size estimates was larger for WT libraries than 3’ libraries. Standard errors trended lower for higher expressed genes with both library types, but there was no significant difference between library types in standard error at any expression level (Figure 2C). However, standard errors were lower for WT libraries for longer transcripts and higher for shorter transcripts (Figure 2D). This corresponded with greater read counts for transcripts with lengths *≈* 1800bp in WT libraries, and lower read counts for shorter transcripts (Figure 2E). Putting this all together we found no average difference in statistical power for discovering differentially expressed genes between WT and 3’ libraries as a function of expression level (Figure 3A). Instead, power was higher for longer genes and lower for shorter genes, as expected (Figure 3B). This means that the distribution of transcript lengths as a function of expression levels determines whether WT or 3’ libraries are likely to discover more differentially expressed genes. For all three species that we studied, WT libraries found 1-5% more differentially expressed genes than 3’ libraries (Table 4), with the advantage of WT over 3’ most evident in herring.

**Figure 2:**
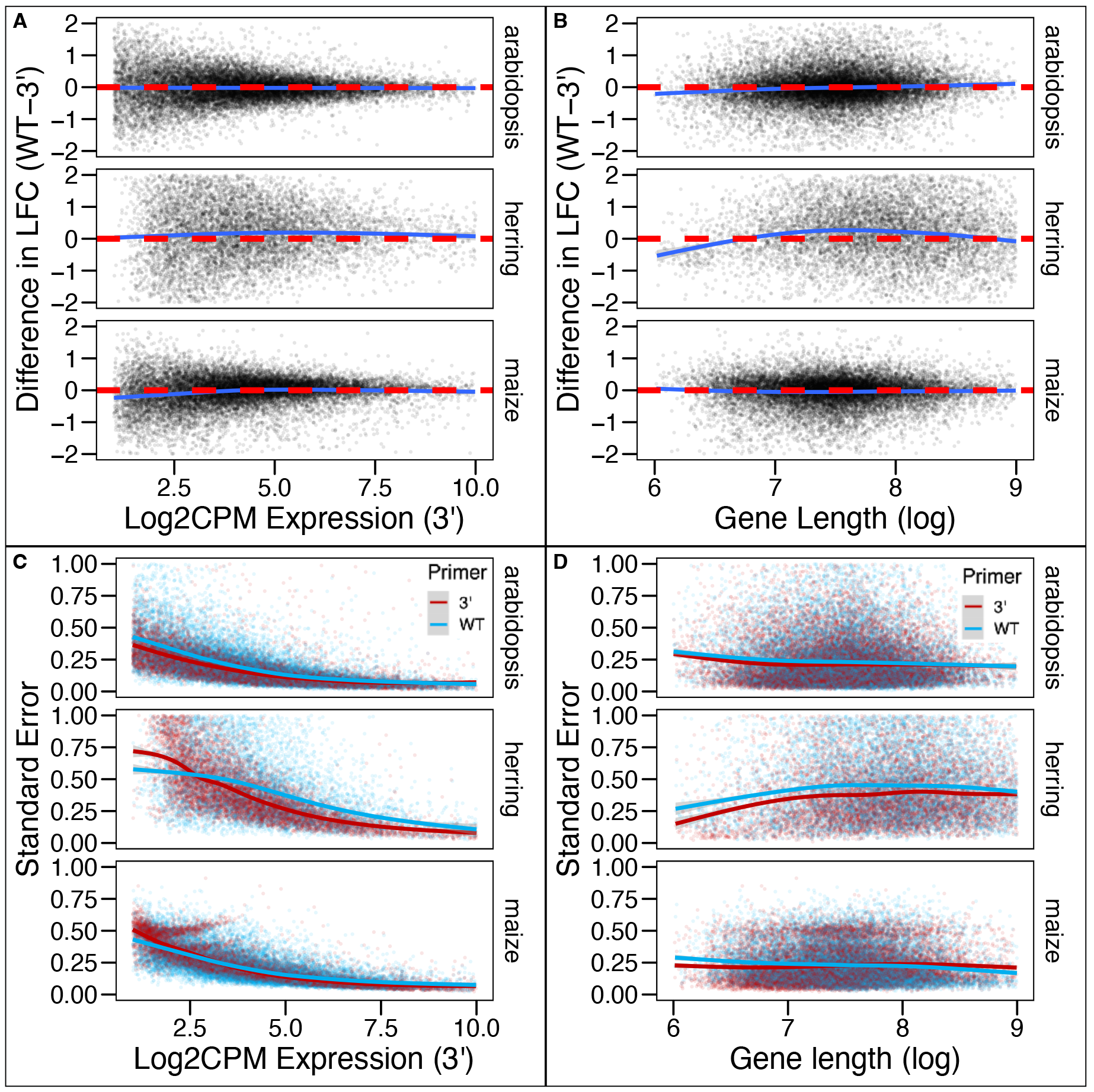
Standard errors but not effect sizes differ between WT and 3’ libraries as a function of gene length but not gene expression. **A**,**B)** Estimated log-foldchanges (LFC) were similar between WT and 3’ libraries across genes as a function of the **A)** gene’s expression (measured as log2 counts per million, log_2_ CPM, by the 3’ library), **B)** or transcript length. Estimated standard errors were similar for the two library types as a function of **C)** expression level, log_2_ CPM), but lower for WT libraries for **D)** longer genes and higher for shorter genes. Each analysis was done separately for each species. Each point in each plot is a single gene, and the smooth curves were generated by the geom smooth function of ggplot2. X and Y-axes were truncated for visualization.

**Figure 3:**
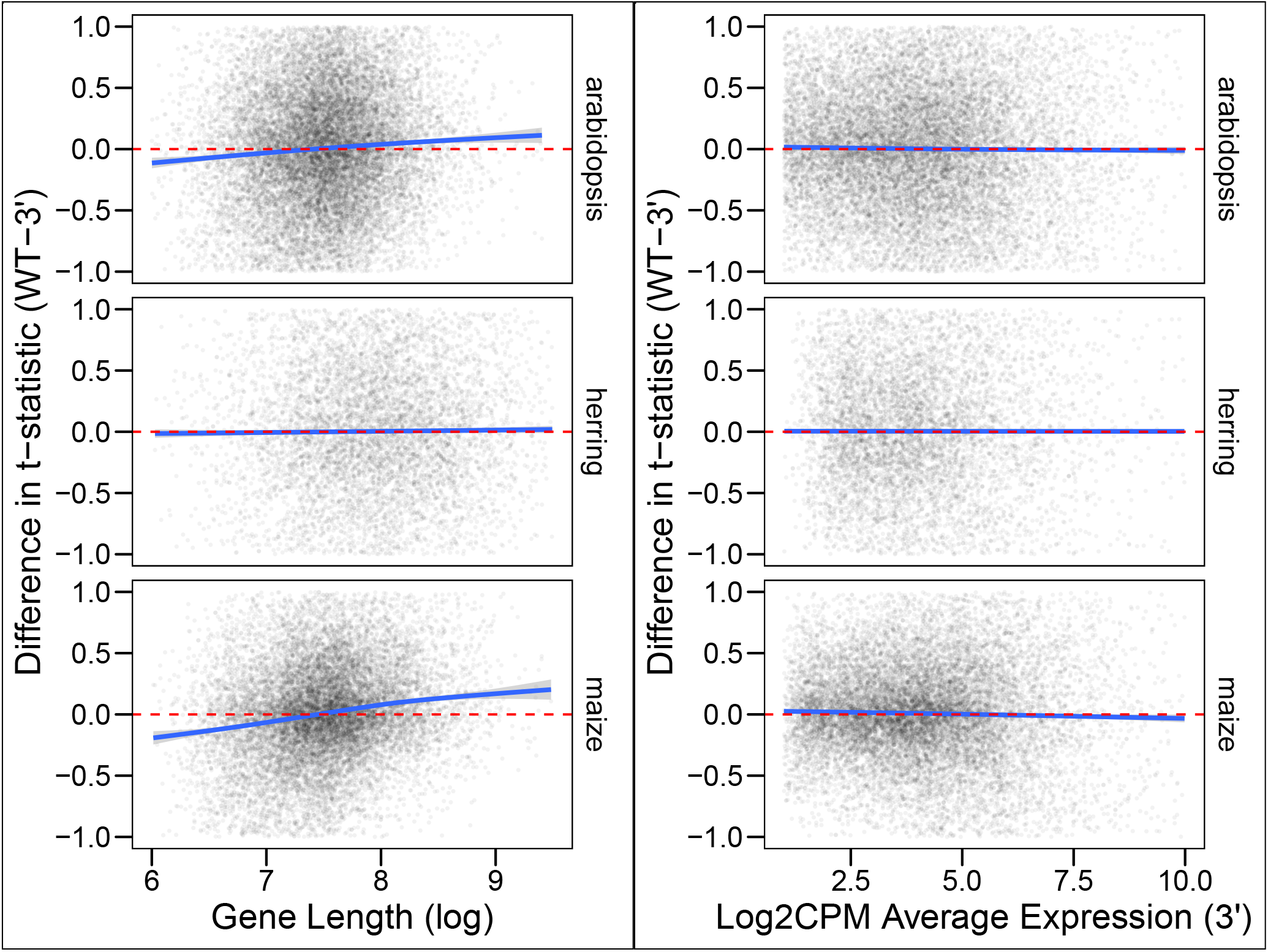
Differential expression analyses of WT libraries have higher power for longer genes, but not for higher-expressed genes than 3’ libraries. **A)** Absolute values of t-statistics were higher on average for longer genes and lower for shorter genes using WT libraries, but did not differ between library priming methods as a function of the **B)** expression of each gene (measured as log2 counts per million, log_2_ CPM, by the 3’ library. Each point in each plot is a single gene, and the smooth curves were generated by the geom smooth function of ggplot2. X and Y-axes were truncated for visualization.

## Discussion

Our analysis and experimental results show that there is no overall benefit of 3’ over WT libraries for expression quantification by RNAseq. WT libraries need not be more expensive than 3’ libraries if comparable kits are available, and WT libraries do not tend to suffer from pseudoreplication or require more sequencing depth to achieve the same precision in expression quantification *on average*. However, length-bias does impact the discovery of differentially expressed genes, such that longer transcripts are more likely to be discovered by WT libraries, and shorter transcripts are more likely to be discovered by 3’ libraries. If researchers knew beforehand which transcripts were of most interest, this could factor into the decision between WT and 3’ libraries. But in general, if all transcripts are equally of interest, there is no clear winner between WT and 3’ for differential expression analyses.

However, WT libraries do have several advantages over 3’ libraries beyond the simple assessment of differential expression at each gene locus. First, mRNA fragments are sequenced from across the entire exonic region of a transcript, which increases the chances of annotating reads (especially in non-traditional model species with low reference genome annotation and assembly quality), as well as alternative splicing and variant detection. The main advantage of 3’ libraries is that they are not subject to the length bias of WT, which can impact downstream analysis such as gene set enrichment [17, 1, 18, 19]. However, expression level also impacts the probability of detecting differential expression for both priming methods, and likely causes a much greater variation in power across genes than variation in length, so this issue is not limited to WT libraries.

Why then have most studies found that WT libraries have more statistical power to detect differentially expressed genes? We also found a similar result, although the difference is small (*≈* 1% *−*5% more genes). The largest difference in numbers of differential gene expression between priming methods in our study was observed in herring. Pacific Herring is the only non-traditional model species in this study. With limited genomic resources, we mapped our RNAseq reads to the publicly available Altantic herring genome assembly. While mapping rates were slightly improved for herring 3’ reads, herring WT libraries quantified over 1,000 additional expressed genes relative to 3’ libraries (Table 3). This may be partially explained by the observation that 3’ UTRs, where priming occurs for 3’ library generation, are less conserved at the sequence level than protein coding sequences [20]. A common alternative for species without a reference is to assemble a transcriptome from the same RNAseq reads used for the differential expression analysis. However, this decision precludes using 3’ priming methods which only retain the 3’ end of gene transcripts [4]. Therefore, WT is a better option for non-traditional model organisms lacking genomic resources. Although further experiments and analysis are necessary to pinpoint the source of increased statistical power inherent in WT libraries, our analysis demystifies the issue of psuedoreplication in WT priming methods and informs a general strategy for applying the wide-ranging utility of RNAseq in biological research.

## Author’s contributions

DER conceptualized the study. TMC and JAG carried out the experiment. TMC and JAG wrote the manuscript with support from DER and AW. All authors contributed critical feedback and helped shape the research analyses and manuscript …

## Acknowledgements

We would like to thank Jane Park for help conceptualizing this work. We would also like to acknowledge Mark Taylor for contributing Arabidopsis tissue …

